# A procedure for solid phase extractions using metal oxide coated silica column in lipidomics

**DOI:** 10.1101/2023.10.15.562428

**Authors:** Hiroaki Takeda, Manami Takeuchi, Mayu Hasegawa, Junki Miyamoto, Hiroshi Tsugawa

## Abstract

Lipid enrichment is indispensable for enhancing the coverage of targeted molecules in mass spectrometry (MS)-based lipidomics studies. In this study, we developed a simple stepwise fractionation method using a titanium- and zirconium-dioxide-coated solid-phase extraction (SPE) silica column that separates neutral lipids, phospholipids, and other lipids, including fatty acids (FAs) and glycolipids. Chloroform was used to dissolve the lipids, and neutral lipids, including steryl esters and di- and triacylglycerols, were collected in the loading fraction. Second, methanol with formic acid (99:1, v/v) was used to retrieve FAs, ceramides, and glycolipids, including glycosylated ceramides and glycosylated diacylglycerols, by competing for affinity with the Lewis acid sites on the metal oxide surface. Finally, phospholipids strongly retained via chemoaffinity interactions were eluted using a solution containing 5% ammonia and high water content (45:50 v/v, 2-propanol:water), which canceled the electrostatic and chelating interactions with the SPE column. High average reproducibility of <10% and coverage of ∼100% compared to those of the non-SPE samples were demonstrated by untargeted lipidomics of human plasma and mouse brain, testis, and feces. The advantage of our procedure was showcased by characterizing minor lipid subclasses, including dihexosylceramides containing very long-chain polyunsaturated FA in the testis, mono- and digalactosyl monoacylglycerols in feces, and acetylated and glycolylated derivatives of gangliosides in the brain that were not detected using conventional solvent extraction methods. Likewise, the value of our method in biology is maximized during glycolipidome profiling in the absence of neutral lipids and phospholipids that cover more than 80% of the chromatographic peaks.

**Table of Contents artwork:** 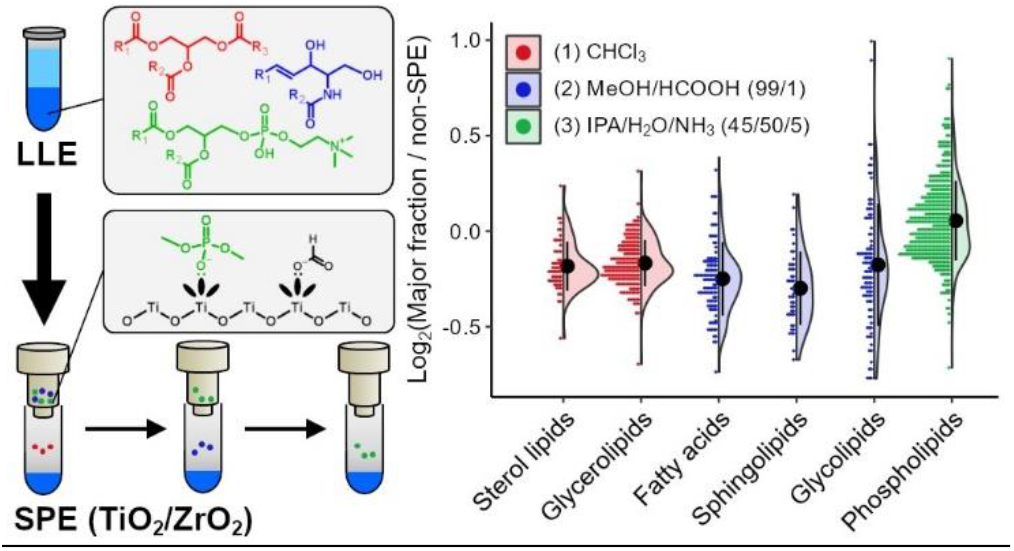

---

Lipids serve as membrane components, signal transduction, and energy storage materials in living organisms. Liquid chromatography coupled with tandem mass spectrometry (LC-MS/MS)-based untargeted lipidomics has made remarkable progress in understanding the diversity of lipids, with approximately 50,000 lipid molecules currently registered in the LIPID MAPS^1^. However, most lipids have a low abundance of MS signals, and low-quality MS/MS spectra increase false hits in the annotation process. This is due to the wide range of lipid concentrations (>10^8^ orders of magnitude in human plasma) of the lipid dynamic range^2^. A small fraction of highly abundant lipid molecules (glycerolipids and phospholipids) obscures the ions of other diverse lipids. This makes it difficult to discover new metabolites that were previously unknown, since high quality MS/MS spectra are preferred to elucidate lipid structures.

This issue may be overcome by improving the fractionation, enrichment, and chromatographic separation to reduce ion suppression, which improves the sensitivity of minor lipids. Preparative LC and thin-layer chromatography are popular techniques for enriching lipids of interest^3,4^, although they are time-consuming processes that need to be optimized for the targeted structure. Alkaline hydrolysis of glycerolipids and phospholipids has been used for the highly sensitive analysis of glycosylated sphingolipids to exploit the difference in bond stability between the amide bond (*N*-acyl chain in sphingolipids) and ester bond (*O*-acyl chain in glycerolipids and phospholipids)^5^. However, the technique is limited to glycosylated sphingolipid analysis, in which the glycerolipids and phospholipids are outside the scope of lipid profiling.

Solid-phase extraction (SPE) is a popular technique for sample preparation, although its application has rarely been reported in lipidomics. Titanium dioxide (TiO_2_) is used to selectively enrich compounds containing a phosphoryl group whose lone pair (Lewis base) interacts with the empty *d* orbital of the TiO_2_ transition metal (Lewis acid)^6^. TiO_2_ is widely used for the enrichment of phosphopeptides^7^ and phospholipids^8,9^. Recently, TiO_2_ has been used as a potential tool for glycolipid fractionation^10,11^. Glycolipids exhibit a high affinity for metal oxide columns via the hydroxyl group of the sugar component under neutral conditions, whereas phospholipid affinity increases under acidic conditions.

Therefore, optimizing the loading and elution solvents when using a TiO_2_ column would facilitate simple and effective fractionation based on polar head categories.

In this study, we developed a procedure to separate lipids into three groups using a commercially available monolithic silica column coated with TiO_2_ and zirconium dioxide (ZrO_2_). Zr has a strong chemical affinity for phosphate groups^12^; thus, glycolipid and phospholipid fractionation are facilitated^13^. The silica phase of the column represents electrostatic interactions with the polar moiety of lipids, helping to enrich glycolipids. We particularly examined the efficiency of our SPE technique for glycerolipids [monoacylglycerols (MGs), diacylglycerols (DGs), and triacylglycerols (TGs)], sphingolipids [ceramides (Cers), hexosylceramides (HexCers), and sphingomyelins (SMs)], and phospholipids [phosphatidylcholines (PCs), phosphatidylethanolamine (PEs), phosphatidylglycerols (PGs), phosphatidylserines (PSs), and phosphatidylinositols (PIs)] by using the authentic standards. Scalability was evaluated using human plasma, mouse brain, testis, and fecal samples.

## EXPERIMENTAL SECTION

### Chemicals and Reagents

Reagent grade ammonia solution (NH_3_) and chloroform (CHCl_3_), ammonium acetate solution for high performance LC (HPLC), formic acid (HCOOH) for LC/MS, acetonitrile (MeCN), methanol (MeOH), 2-propanol (IPA), and ultrapure water (H_2_O) for quadrupole time-of-flight MS (QTOFMS) were obtained from FUJIFILM Wako Pure Chemical Corp. (Osaka, Japan). Toluene for HPLC was obtained from Sigma-Aldrich (Tokyo, Japan). MonoSpin Phospholipid was purchased from GL Sciences Inc. (Tokyo, Japan) for the monolithic silica column coated TiO_2_and ZrO_2_ (*φ* 4.2 μm × 1.5 mm). Synthetic standards and EquiSPLASH containing 100 μg/mL of deuterium-labeled standards were purchased from Avanti Polar Lipids Inc. (Alabaster, AL, U.S.A.). Human plasma (pooled in heparin sodium) was obtained from Cosmo Bio Co. Ltd. (Tokyo, Japan).

### Animals

All animal experiments were performed in accordance with the ethical protocol approved by Tokyo University of Agriculture and Technology (R5-50). C57BL/6J male mice were purchased from SLC (Shizuoka, Japan). The mice were fed the chow of CE-2 (CLEA Japan, Tokyo, Japan). The brain, testis, and feces were harvested, immediately frozen after dissection, and stored at **−**80°C until extraction.

### Monophasic liquid extraction

Monophasic lipid extraction was performed as described previously^14^. Samples were mixed with 200 μL of MeOH and extracted using ultrasonication for 5 min. One hundred microliters of CHCl_3_ was added and the solution was vortexed for 1 min and extracted on ice for 60 min. The solution was further mixed with 20 μL of H_2_O and vortexed for 1 min. The supernatant was collected after centrifugation at 16,000 g for 5 min at 4°C. Lipid extracts were dried using a centrifuge evaporator.

### Biphasic liquid extraction

The Bligh and Dyer method was performed according to a previously described method with minor modifications^15^. Samples were mixed with 1,000 μL of MeOH/CHCl_3_/H_2_O (10:4:4, v/v/v). Lipids were extracted using a vortex mixer for 1 min and then ultrasonicated for 5 min. The solution was centrifuged at 16,000 ×*g* for 5 min at 4°C, and 700 μL of the supernatant was transferred to a clean tube. The supernatant was mixed with 235 μL of CHCl_3_ and 155 μL of H_2_O using a vortex mixer for 1 min. Both layers were collected after centrifugation at 16,000 ×*g* for 5 min at 4°C. Lipid extracts were dried using a centrifuge evaporator.

### Solid phase enrichment

A dried lipid extract was dissolved by applying 100 μL of loading solvent, and 50 μL of the sample was loaded to the SPE column. An equal amount of EquiSPLASH was added to each sample before the purification of the lipid extract. The detailed protocol developed in the present study is described in the Supporting Information. All fractions collected via SPE were dried using a centrifugal evaporator and dissolved in MeOH containing Cer 18:1;O2/17:0 standard for LC-MS analysis.

### Lipidome analysis

The LC/QTOFMS system comprised an Exion LC and ZenoTOF 7600 with an electrospray ionization ion source (SCIEX, Framingham, MA, U.S.A.). The LC conditions were as follows: injection volume, 1 μL; mobile phase, MeCN/MeOH/H_2_O (1:1:3, v/v/v) (A) and MeCN/IPA (1:9, v/v) (B) (both contained 5 mM of ammonium acetate and 10 nM of ethylenediaminetetraacetic acid); flow rate, 300 μL/min; column; Unison UK-C18 MF (50 × 2.0 mm, 3 μm, Imtakt Corp., Kyoto, Japan); gradient, 0.5% (B) (1 min), 0.5− 40% (B) (4 min), 40−64% (B) (2.5 min), 64−71% (B) (4.5 min), 71−82.5% (B) (0.5 min), 82.5−85% (B) (6.5 min), 85− 99% (B) (1 min), 99% (B) (2 min), 99−0.5% (B) (0.1 min), 0.5% (B) (2.9 min); column oven temperature, 45°C. The MS conditions were as follows: ion source gas 1, 50 psi; ion source gas 2, 50 psi; curtain gas, 35 psi; CAD gas, 7; source temperature, 300°C; spray voltage, 5500 V (positive) or **−**4500 V (negative); *m/z* range of TOF-MS, 75−1250 Da (positive) or 100−1700 Da (negative); declustering potential, **−**80 V; collision energy, 42 ± 15 eV; Q1 resolution, unit; acquisition mode, data dependent MS/MS; accumulation time, 50 msec.

### Data analysis

The lipid molecules were annotated using MS-DIAL version 5.1^14^. The data processing parameters and the adduct ions used to calculate lipid amounts are listed in Tables S1 and S2. The annotated results were curated manually. The data matrix was prepared using the mean value of the analytical replicates (*n* = 4) after the normalization of the lipid molecules based on the sample volume and ion abundance of the Cer 18:1;O2/17:0 standard (Table S3). The lipidome results are summarized in Tables S4–10.

## RESULTS AND DISCUSSION

### Optimization of the fractionation protocol

Lewis acidic sites on the metal oxide surface have an affinity for lone-pair heteroatoms. Seven solutions including hexane, toluene, CHCl_3_, MeOH, MeOH/HCOOH (99/1, v/v), MeOH/H_2_O/NH_3_ (80/15/5, v/v/v), and IPA/H_2_O/NH_3_ (45/50/5, v/v/v) were investigated to determine the lipid solubility and interaction with stationary phases by using mouse brain extract and synthetic standards. Hexane, toluene, and CHCl_3_ are commonly used in LC-based lipidomics and exhibit sufficient solubility for a wide range of lipid subclasses (Figure S1). Although MeCN was a suitable solvent for the adsorption of lipids (except for glycerolipids) on a TiO_2_ column in a previous report ^10^, the solubility of lipids was insufficient when MeCN was used in our study (except for DG, TG, Cer, fatty acids (FA), PC, and PG). The optimal solvent for loading the lipid extract was determined based on solubility. Lipid molecules primarily bind to the stationary phase of silica-based columns via electrostatic interactions. The elution behavior of steryl esters (SEs), TGs, DGs, Cers, and glucosylceramides (GlcCers) depends on solvent polarity (hexane, toluene, CHCl_3_, and MeOH) (Figure 1a). We selected CHCl_3_ as a suitable solvent to dissolve lipids and elute the major neutral lipids of DGs, TGs, and SEs, while retaining sphingolipids and phospholipids.

**Figure 1.**
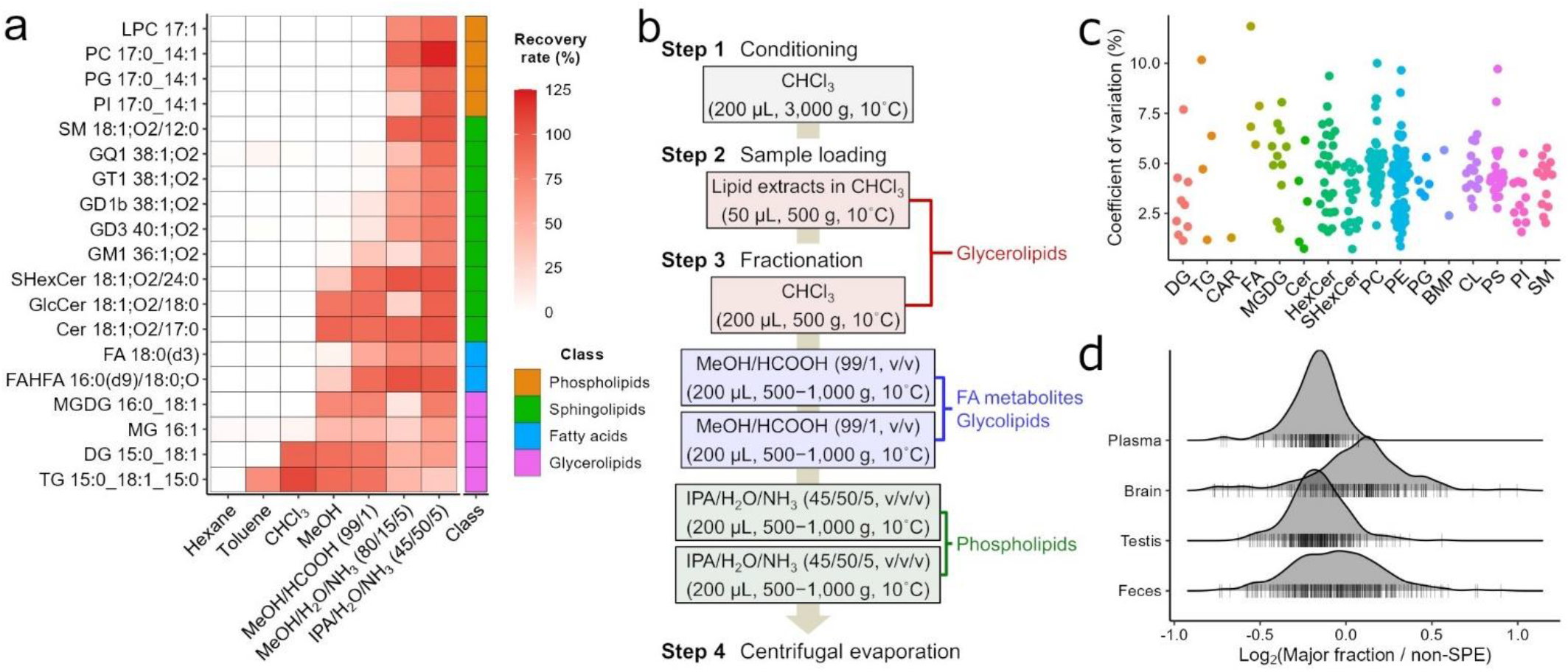
Development of the stepwise fractionation. (a) Recovery rate of the loading fraction obtained by dissolving the lipid standards in each solvent. (b) Workflow of the SPE-based fractionation. (c) Evaluation of repeatability using the mouse brain extract. The coefficient of variation was evaluated using the SPE replicates (*n* = 4). In addition to the subclasses confirmed by authentic standards, acyl-carnitine (CAR), bis(monoacylglycero)phosphate (BMP) and cardiolipin (CL) were evaluated. (d) Density plot of four biological samples. Relative abundance was calculated by comparing lipidomics data from non-SPE samples with the major fractions obtained by SPE. The vertical markers represent the ratio of each molecular abundance between SPE and non-SPE.

Secondly, the fractionation of lipid groups with hydroxyl and/or carboxyl groups, such as FAs, sphingolipids, and glycolipids, was investigated. Among the three functional groups (phosphate, hydroxyl, and carboxyl), the phosphate group has the strongest chemical affinity for metal oxides. The addition of HCOOH affects the competition with organic ligands because HCOOH has a carboxyl group that has a chemical affinity for metal oxides. Compared to MeOH without HCOOH, MeOH with HCOOH effectively eluted FAs, FA esters of hydroxy FAs (FAHFAs), sulfatides (SHexCers), and some gangliosides (such as the GM and GD series) with high recoveries (Figure 1a and S2). A previous study reported that acidic solvents keep a strong affinity for phospholipids and weak interactions with glycolipids when using a TiO_2_ column^11^. Therefore, MeOH/HCOOH (99/1, v/v) was found to be the optimal solvent for eluting FA metabolites, Cers, and glycolipids, including monogalactosyl DGs (MGDGs), HexCers, and SHexCers. The HCOOH can be removed using a centrifugal evaporator. The throughput and cost of purification were reduced by avoiding the removal step for counter substances (such as 2,5-dihydroxybenzoic acid^13^).

The phosphate group has strong Lewis basic sites. Therefore, phospholipid elution can be achieved using a solvent containing NH_3_ that removes chelating interactions. Most phospholipids were eluted using MeOH/H_2_O/NH_3_ (80/15/5, v/v/v), whereas PIs with six hydroxyl groups in the inositol moiety were retained because of their strong affinity for metal oxide columns. Electrostatic and chelating interactions are canceled by the hydration of polar groups and ammonium ions, respectively, to elute lipids with rich polar functional groups. Therefore, a solvent with a high H_2_O content was used. Lipid solubility was maintained by changing the solvent from MeOH to IPA (Figure S1a). Finally, IPA/H_2_O/NH_3_ (45/50/5, v/v/v) was determined as the optimal solution to achieve a high recovery rate (Figure 1a).

We proposed a procedure for the stepwise purification of lipid subclasses (Figure 1b and Supporting Information). The repeatability of the protocol was within 10% in the lower phase of the mouse brain extract using the Bligh and Dyer method (Figure 1c). Meanwhile, solvents mixed with NH_3_ caused the acyl migration of lysophospholipids and slight alkaline hydrolysis effect of phospholipids (less than 1%) (Figure S1b and S1c). Because the quantitative reliability of lysophospholipids is compromised when samples contain high amounts of phospholipids and/or *sn*-2 positional isomers of lysophospholipids, we believe that lysophospholipids are out of the scope of our SPE-based lipidomics study.

### Application to untargeted lipidomics

The optimized SPE procedure was applied to human plasma, mouse brain, testis, and feces to evaluate the performance of SPE with respect to a wide variety of lipid subclasses. The sample volume of each extraction protocol injected into the column was kept constant to evaluate the degree of ion suppression and the recovery rate of each lipid extract. In the present study, the differences among plasma, brain, testis, and fecal samples were investigated using monophasic extraction, where unwanted salts and protein residues can be excluded via the SPE procedure. The lipid distributions among the four samples were similar when evaluating the relative abundance between the non-SPE and the main fraction obtained by SPE (Figure 1d). Lipids were fractionated with high recovery into three optimized solvents, despite differences in the sample matrix (Figure 2a). This is consistent with the synthetic standards. Minor glycolipids, which were outside the dynamic range owing to abundant lipids, were detected in mouse testis and feces owing to the removal of neutral lipids and phospholipids. In the testis, the amount of lipids in the acidic MeOH fraction was approximately 50-fold lower than that in the other fractions (Figure 2b). Previous studies have reported that very long-chain polyunsaturated FAs (VLC-PUFAs) with more than 24 carbons are found in different lipid classes such as SM, Cer, SE, and TG^16,17^. The SPE method enabled us to detect 41 sphingolipids containing VLC-PUFA, including HexCer and Hex2Cer, which have rarely been reported in previous studies (Figure 2c and 2d and Table S8). Moreover, fecal analysis detected MGDG and digalactosyl DG (DGDG) before and after sample purification, whereas monogalactosyl MG (MGMG) and digalactosyl MG (DGMG) were only detected in the acidic MeOH fraction (Figure 2e and 2f). Likewise, the stepwise fractionation should be an important tool for the exploratory study of glycolipidome since the removal of neutral lipids and phospholipids using SPE decrease ion suppression in MS analysis (Figure S6b**−**6f)^18^. Although we believe that the use of a C18-SPE column is the well-validated method for purifying gangliosides, our SPE method was used to enrich the brain gangliosides. A total of 61 gangliosides, including acetylated and glycolylated derivatives as previously reported^19,20^, were fractionated simultaneously by collecting IPA/H_2_O/NH_3_ (45/50/5, v/v/v) immediately after loading the upper phase of the Bligh and Dyer method (Figure 2g, 2h, and Table S9), while they were not within the scope of the MS-DIAL lipidomics pipeline^14^. Since the fractionation of gangliosides depends on the number of sialic acids with Lewis base sites (Figure 1a), detailed fractionation based on their polar groups, such as GM, GD, GT, and GQ, would become possible by further optimizing our SPE procedure in the future.

**Figure 2.**
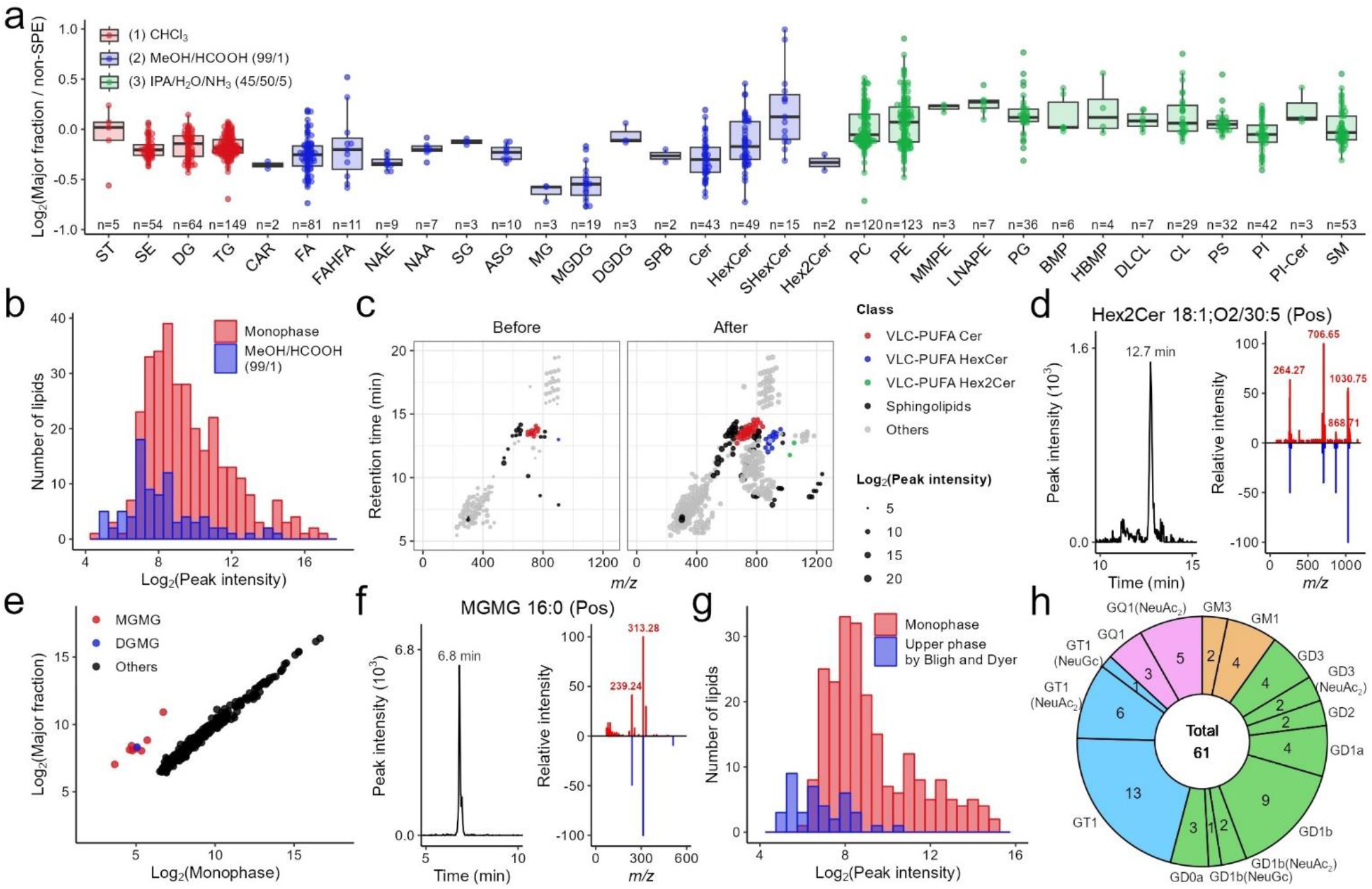
Lipidome analysis using the fractionation method. (a) Lipidome analysis of each fraction. Lipidomics data from four biological samples were combined by lipid subclass. Sterol (ST), *N*-acyl-ethanolamine (NAE), *N*-acyl amide (NAA), sterylglucoside (SG), acyl-SG (ASG), sphingoid base (SPB), monomethyl-PE (MMPE), *N*-acyl-lyso-PE (LNAPE), Hemi BMP (HBMP) and dilyso-CL (DLCL) were also grasped by the SPE method. (b) Distribution of lipid abundances in the mouse testis extract. (c) Testicular lipidome analysis in the acidic MeOH fraction before and after enrichment (each, 0.5- and 1.3 × 10^2^ μg dried weight in the liquid injected into the column, respectively). (d) Extracted ion chromatogram and the MS/MS spectrum pattern of Hex2Cer which was detected by the enriched sample of testis lipid extract. (e) Comparison of lipidomics data between with or without SPE obtained from the mouse feces. (f) An example of unique molecules only detected in the acidic MeOH fraction of mouse feces. (g) Distribution of lipid abundances in the mouse brain extract. (h) Number of gangliosides detected in the mouse brain. Three sialic acids [*N*-acetylneuraminic acid (NeuAc), NeuAc_2_, and *N*-glycolylneuraminic acid (NeuGc)] which are absent in MS-DIAL program were detected in this study.

## CONCLUSION

We developed a simple and effective lipid separation method using TiO_2_ and ZrO_2_ coated silica column, wherein three lipid categories (neutral lipids, phospholipids, and other lipids, including FA metabolites and glycolipids) were fractionated, while SPE is often used to divide a mixture into only two categories. In addition, the recovery rate of lipid molecules in the plasma, brain, testis, and fecal samples was 80–120%. This stepwise solid-phase fractionation contributes to reducing false positive discoveries in untargeted lipidomics by combining the knowledge of separation behavior based on functional groups. In addition, the SPE method at the microscale can accelerate the structural analysis of novel compounds using advanced fragmentation techniques, such as electron activated dissociation^21^. There are few reports on high-recovery glycolipid fractionation. Therefore, this technique may facilitate the discovery of novel glycolipid molecules in biological samples, as demonstrated by the characterizations of HexCer and Hex2Cer containing VLC-PUFAs in the testis, MGMG and DGMG in feces, and ganglioside derivatives in the brain. The untargeted lipidomics of an enriched fraction, named untargeted “probing” lipidomics facilitates the discovery of novel lipid metabolites associated with phenotypes. Furthermore, lipidomics using low-flow LC (<10 μL/min), wherein a simplified sample matrix is preferable for a sustainable operation, can be accelerated by using our fractionation procedure.

## ASSOCIATED CONTENT

### Supporting Information

The Supporting Information is available free of charge at ACS Publications website.

### Data availability

All raw MS data are available on the RIKEN DROPMet website (http://prime.psc.riken.jp/menta.cgi/prime/drop_index) under index number DM0051. The MS-DIAL source code is available at https://github.com/systemsomicslab/MsdialWorkbench.

## AUTHOR INFORMATION

### Author Contributions

Hiroaki T. and Hiroshi T. designed the study. M.H. and J.M. prepared the animals. Hiroaki T. and M.T. extracted the samples, analyzed the lipids, and performed the data analysis. Hiroaki T. and Hiroshi T. wrote the manuscript and prepared the figures. All the authors approved the final version of the manuscript.

## ACKNOWLEDGMENT

The authors thank Dr. Masahiro Furuno (Osaka University) and Dr. Shigenori Ota (GL Sciences, Inc.) for technical advice on chemoaffinity. This research was supported by AMED Brain/MINDS (JP15dm0207001, Hiroshi T.), JST ERATO (JPMJER2101, Hiroshi T.), the JSPS KAKENHI (21K18216, Hiroshi T.), National Cancer Center Research and Development Fund (2020-A-9, Hiroshi, T.), AMED Japan Program for Infectious Diseases Research and Infrastructure (21wm0325036h0001, Hiroshi T.), and Technologically Advanced research through Marriage of Agriculture and engineering as Groundbreaking Organization (TAMAGO to J.M. and Hiroshi T.).

