## Supplementary material for "A procedure for solid phase extractions using metal oxide coated silica column in lipidomics": Supporting_Information.pdf

**Contents**

Supplements of Experimental Section

Supplementary Figure 1–6

Supplementary Table 1–10

**Optimized SPE protocol for untargeted lipidomics.** MonoSpin Phospholipid (GL Sciences) was used for the SPE. A dried lipid extract was dissolved by adding 100 µL of $CHCl_3$, to which an equal amount of EquiSPLASH was added. The SPE columns were conditioned with 200 µL of $CHCl_3$ before loading the lipid extract. The lipid extract in $CHCl_3$ (50 µL) was applied to the SPE columns and then centrifuged at 500 ×*g* for 2 min at 10°C to elute neutral lipids. To increase the recovery of neutral lipids, 200 µL of $CHCl_3$ was added to the SPE columns and collected in the same tube after the centrifugation. Secondly, 200 µL of MeOH/HCOOH (99:1, /v/v) was applied to the SPE columns and then centrifuged at 500 ×*g* for 5 min at 10°C to elute FA metabolites, Cers, and glycolipids. The fraction was also collected twice to increase the recovery. Finally, 200 µL of IPA/$H_2O$/$NH_3$ (45/50/5, v/v/v) was applied to the SPE columns and then centrifuged at 500 ×*g* for 5 min at 10°C to elute phospholipids. Phospholipids were eluted in duplicate for high recovery. If the samples did not complete elution under the centrifugation conditions, the SPE columns were centrifuged at 1000 ×*g* for 5 min at 10°C. The samples were dissolved in MeOH containing Cer 18:1;O2/17:0 standards after drying in a centrifugal evaporator.

**Lipid analysis.** The total ion chromatograms for each sample are shown in Figure S3. The peak shapes of acidic lipids such as PS were improved by adding 10 nM of EDTA to both mobile phases. In a previous report, EDTA improved peak shape and sensitivity of acidic phospholipids such as lysophosphatidic acid, phosphatidic acid, and PS by pre-washing the column with EDTA and adding it to the sample solution [1]. EDTA suppresses interactions between the column materials and acidic moieties of lipids by forming complexes with metal ions. In the present study, EDTA was added to the mobile phase to improve the chromatographic peak shapes of the PS molecules, because our LC system included steel use stainless tubes and fittings.

**Data normalization.** All the samples were dissolved in MeOH containing Cer 18:1;O2/17:0 standards for LC-MS analysis. Although the recovery rate of each lipid class was within a reasonable range using synthetic standards (Figure 1a), the relative abundance of some lipids, such as PC and PE, before and after lipid purification exceeded 200% in the negative ion mode (Figure S4a–4d). The molecules included PC 16:0_18:2 (10.1 min), PC 16:0_20:4 (10.1 min), PC 18:1_20:4 (10.2 min), and PE 16:0_22:6 (10.2 min) in the brain, and PC 16:0_18:2 (10.1 min) and PE 15:0_16:0 (10.2 min) in the feces, where the retention times are described in parentheses (Tables S5 and S6). This indicates that all lipids were detected within the same retention time frame (10.1–10.2 min). For example, PC 15:0_18:1(d7) elutes at 10.2 min using an analytical method, which may result in inappropriate normalization when EquiSPLASH is used for quantification. In contrast, Cer 18:1;O2/17:0 showed a constant response (11.87%) in the plasma, brain, testis, and feces, with or without the SPE procedure (Figure S4e). This suggests that the other molecules did not interfere with the ionization of Cer 18:1;O2/17:0. Although the normalization of Cer 18:1;O2/17:0 cannot be used to calculate the concentration of each lipid molecule, it is adequate to normalize the analytical error. Therefore, the type of quantification corresponds to level 3 (lipid class other than analyte or no co-ionization of analyte and internal standard) of the Lipidomics Standards Initiative International Guideline [2]. The differences in the normalization of lipid recovery between EquiSPLASH and Cer 18:1;O2/17:0 are summarized in Figure S5.

We prepared the data matrix using the following procedure, based on the characteristics of the current RPLC/MS system. First, lipidomics data were normalized using peak intensity of the Cer 18:1;O2/17:0 standard. Second, the average of the normalized values among the technical replicates ($n$ = 4) was calculated for all lipids. Third, the average values were normalized to the sample volume injected into the column. Finally, the relative abundance was calculated as follows:

$$Relative\ abundance\ of\ lipid\ X = \frac{The\ average\ value\ of\ lipid\ X\ with\ SPE}{The\ average\ value\ of\ lipid\ X\ without\ SPE} \times 100\ (\%)$$

The average value of a lipid "with SPE" is calculated from the lipidomics data of a major fraction for the lipid.

**Difference in lipidomics data between monophasic and biphasic extractions.** The sample volume of each extraction protocol injected into the column was kept constant to evaluate the degree of ion suppression and the recovery rate of each lipid extract. There was no significant difference between the two extractions, except for SE and TG, in the mouse testis using the current reversed-phase LC/MS system (Figure S6a). SE and TG levels increased with increasing lipid content in the plasma samples when monophasic extraction was used for lipid extraction [3]. The same trend was observed for testicular lipidome analysis only, although the difference in sample weight between the brain and testicular lipidome analyses was within two-fold in the present study.

### Supplementary Figures

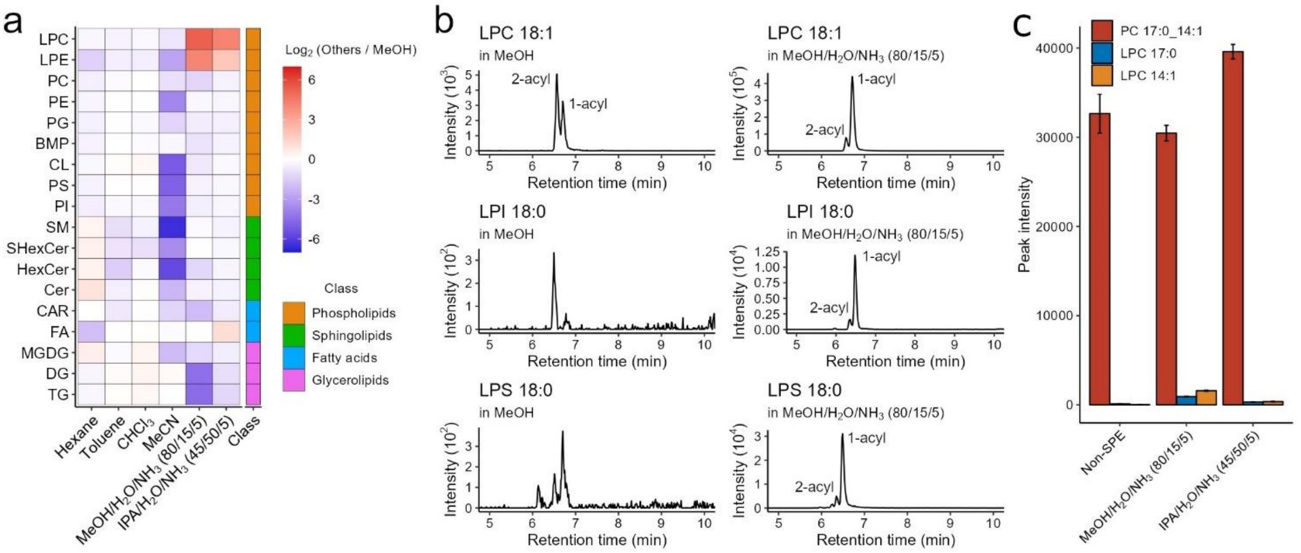


**Figure S1. Optimization of loading and elution solvents for lipid enrichment.** (**a**) Solubility of lipid molecules in each solvent. Dried lipid extracts of mouse brain were dissolved in each solvent after evaporation. Values were calculated using the mean value of the $log_2$- transformed relative intensity (each solvent vs. MeOH) per lipid class. (**b**) Evaluation of acyl migration of lysophospholipids and alkaline hydrolysis of phospholipids using alkaline solvent. The lower phase of the mouse brain extract was dissolved with MeOH or MeOH/$H_2O$/$NH_3$ (80/15/5, v/v/v). (**c**) Examination of the degree of degradation products using the synthetic standard of PC 17:0_14:1.

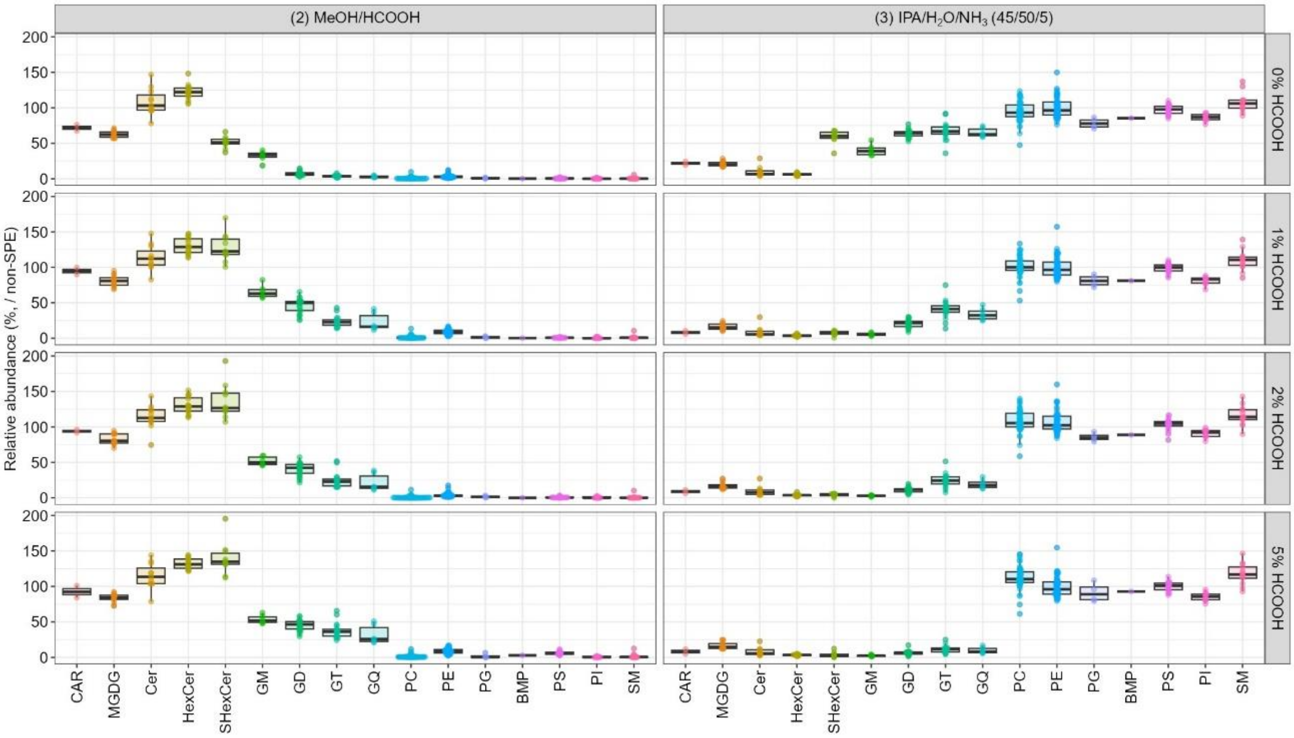


**Figure S2. Change in lipid recovery in the mouse brain extract by adding HCOOH.** Lipids were extracted by the Bligh and Dyer method. Relative abundance on the vertical axis was evaluated by comparison with lipidomics data from non-SPE samples.

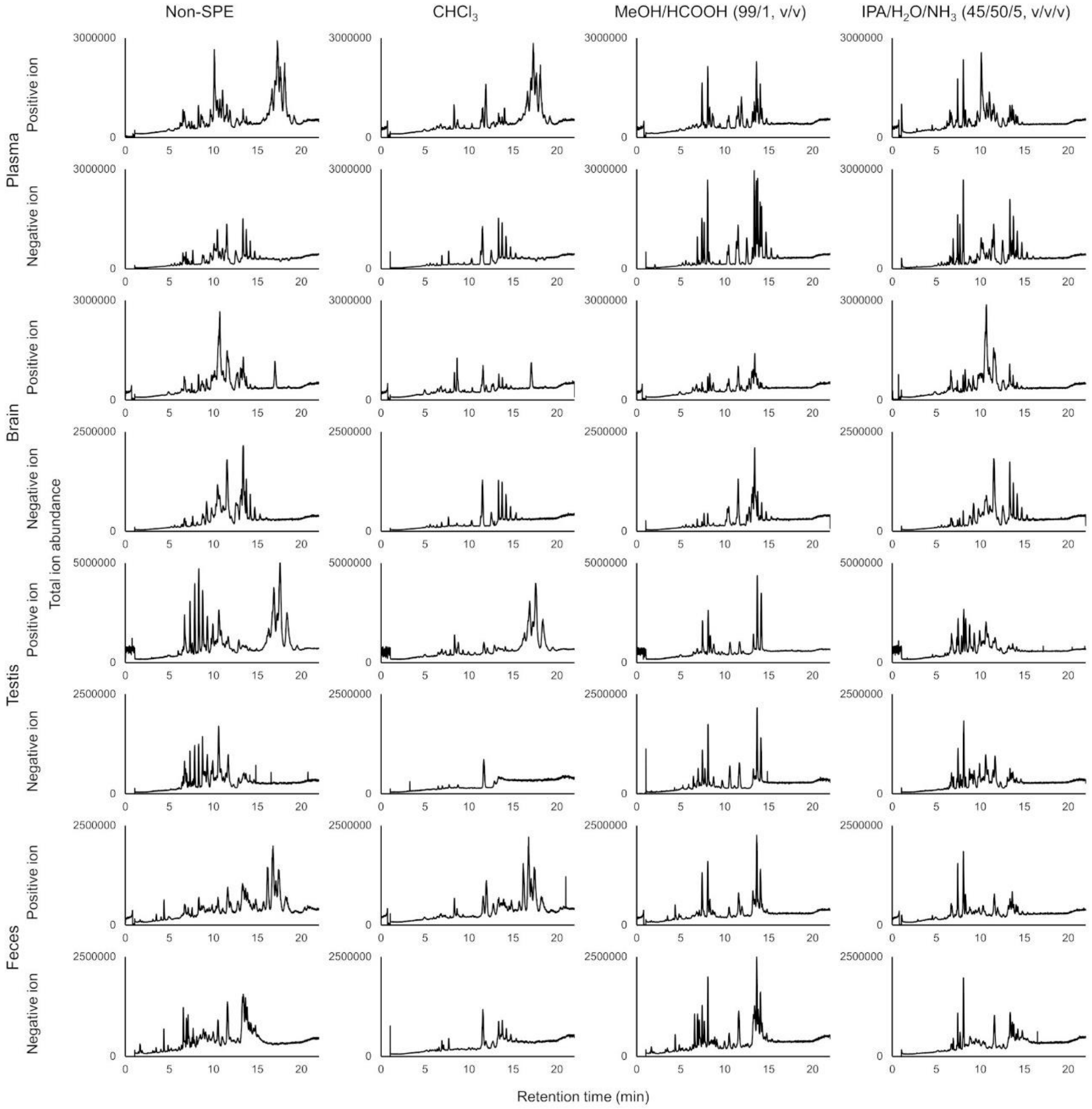


**Figure S3. Total ion chromatogram for each sample.**

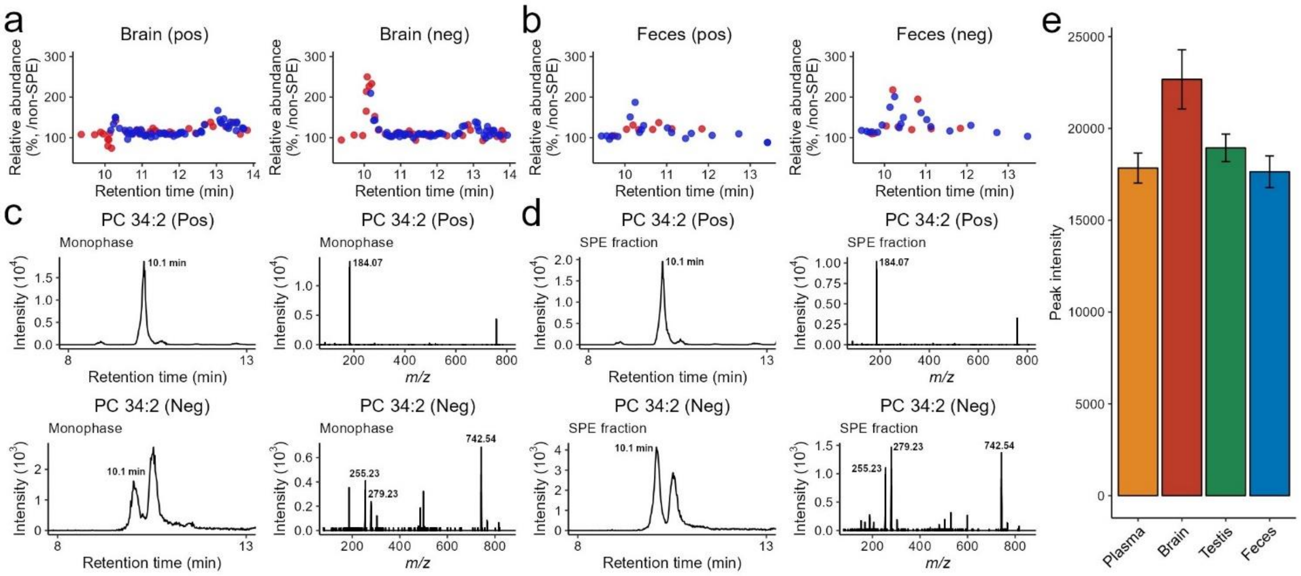


**Figure S4. Ion suppression of lipids at specific retention times.** (**a**, **b**) Relative abundances of PC and PE in mouse brain (**a**) and feces (**b**) in both positive and negative ion modes. Relative abundances on the vertical axis were evaluated by comparison with lipidomics data from non-SPE samples. The red and blue dots represent the molecules of PC and PE, respectively. (**c**, **d**) Example of chromatogram and MS/MS spectrum of brain lipids with a relative abundance > 200% before (**c**) and after (**d**) the SPE procedure. (**e**) Peak intensity of Cer 18:1;O2/17:0 standard in each biological sample.

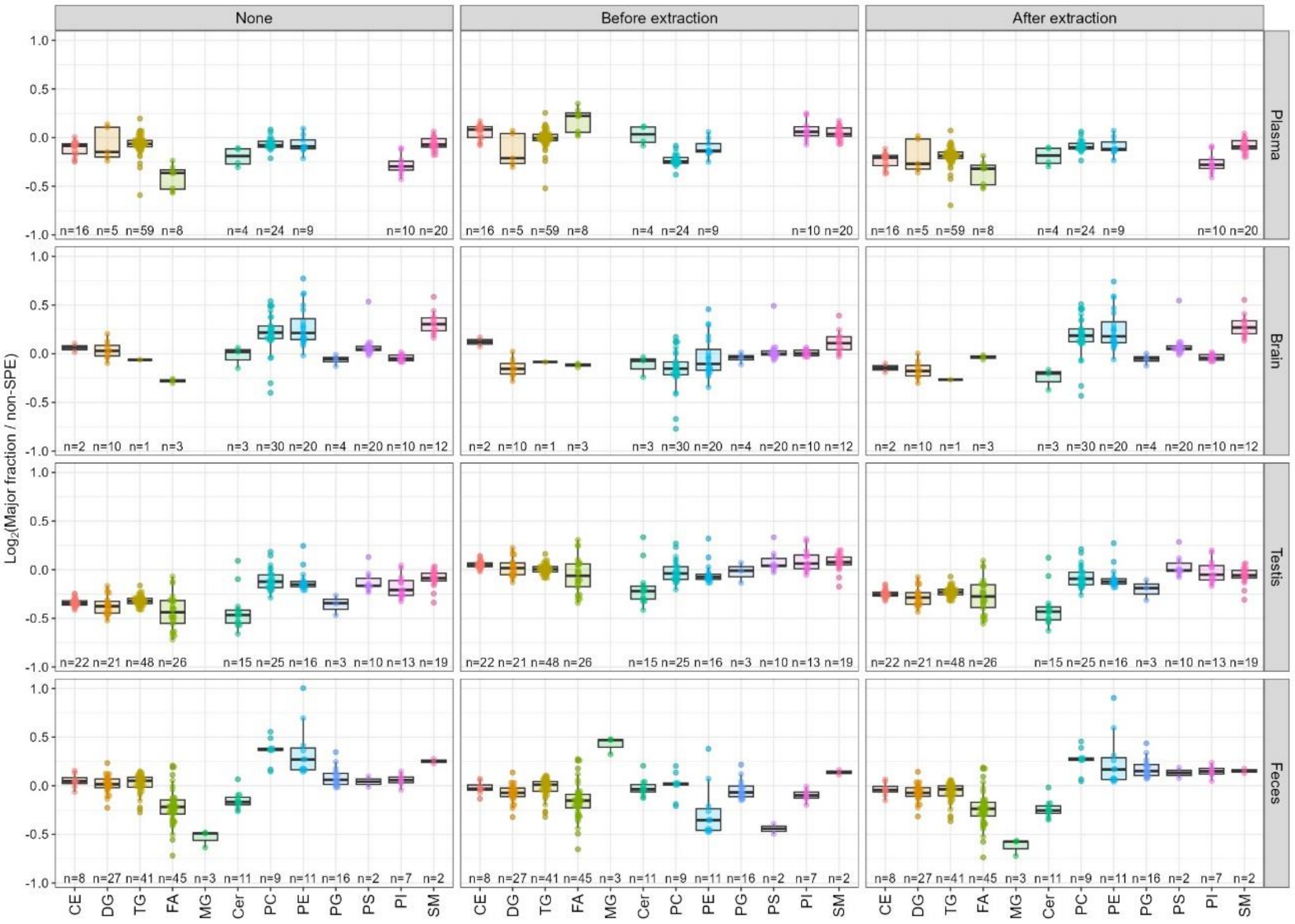


**Figure S5. Normalization of lipid recovery using internal standards.** Lipidomics data in the middle panel was normalized using EquiSPLASH and a deuterium labeled FA mixture (FA 16:0(d3) and FA 18:0(d3)) that was added before the SPE procedure (10 nL of undiluted solution and 1 pmol of FAs were injected into the column, respectively). Lipidomics data in the right panel was normalized using Cer 18:1;O2/17:0 that was added after the SPE procedure (2 pmol injected into the column).

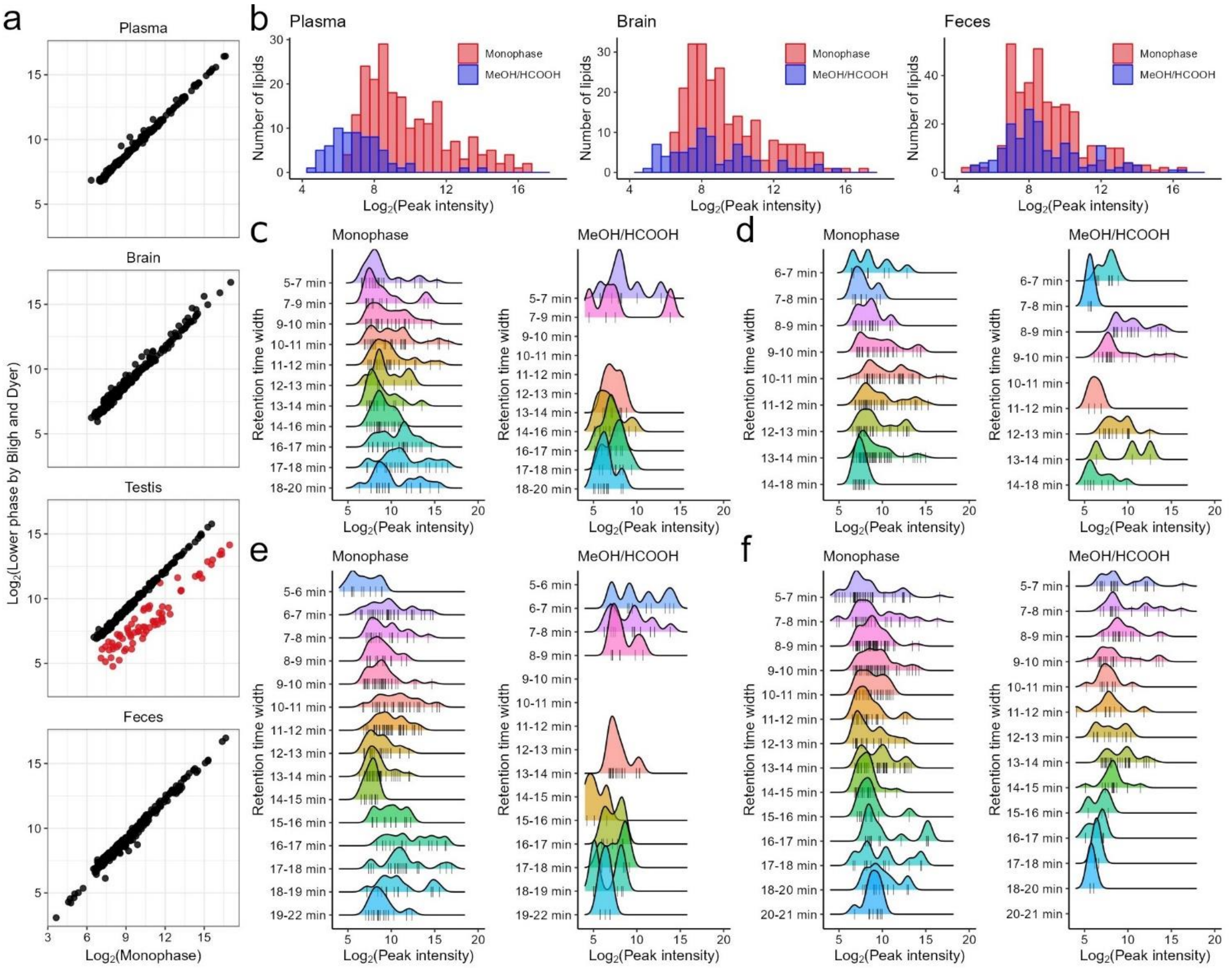


**Figure S6. Lipidome analysis of biological samples.** (**a**) Comparison of extraction protocols between monophasic and biphasic liquid extractions. The sample volume of each extraction protocol injected into the column was kept constant to evaluate the degree of ion suppression and the recovery rate of each lipid extract. The red dots represent SE and TG. (**b**) Lipid distribution of plasma, brain, and feces before and after the SPE procedure. The lipid distribution of testis is shown in Figure 2b. (**c**–**f**) Ridgeline plot of plasma (**c**), brain (**d**), testis (**e**), and feces (**f**) under each RT range. The rug plot represents the molecules.

### Supplementary Tables

**Table S1. Parameters of MS-DIAL software.**

**Table S2. Summary of lipid subclasses and their adduct ions used to calculate lipid amounts in the study.**

**Table S3. Amount of biological samples injected into the column.** [a] Lyophilized volume of mouse brain and feces injected into the column. [b] FA 16:0(d3) and FA 18:0(d3) were added before SPE. [c] Cer 18:1;O2/17:0 was added after SPE.

**Table S4. Lipidome analysis of human plasma.** Lipidomics data were normalized using the sample volume (0.02 μL) and Cer 18:1;O2/17:0 added after the SPE procedure (2 pmol).

**Table S5. Lipidome analysis of mouse brain.** Lipidomics data were normalized using the sample volume ($3.0 \times 10^{-4}$ mg dry weight) and Cer 18:1;O2/17:0 added after the SPE procedure (2 pmol).

**Table S6. Lipidome analysis of mouse testis.** Lipidomics data were normalized using the sample volume ($5.0 \times 10^{-4}$ mg dry weight) and Cer 18:1;O2/17:0 added after the SPE procedure (2 pmol).

**Table S7. Lipidome analysis of mouse feces.** Lipidomics data were normalized using the sample volume ($4.0 \times 10^{-3}$ mg dry weight) and Cer 18:1;O2/17:0 added after the SPE procedure (2 pmol). NAGly, *N*-acyl glycine; NAGlySer, *N*-acyl glycyl serine; NAOrn, *N*-acyl ornithine.

**Table S8. Detailed lipid profiling of the enriched acidic MeOH fraction in mouse testis.** Lipidomics data were normalized using the sample volume ($1.3 \times 10^{-1}$ mg dry weight) and Cer 18:1;O2/17:0 added after the SPE procedure (2 pmol). NATau, *N*-acyl taurine.

**Table S9. Lipid profiling of upper phases collected using the Bligh and Dyer method in mouse brain.** Lipidomics data were normalized using the sample volume ($2.0 \times 10^{-2}$ mg dry weight) and Cer 18:1;O2/17:0 added after the SPE procedure (2 pmol).

**Table S10. Change in lipid recovery in the mouse brain extract following addition of HCOOH.**
